# Comparative Analysis of Neonatal Hypomyelination Models Using Spatial Transcriptomics

**DOI:** 10.1101/2025.06.23.661209

**Authors:** Rafael Bandeira Fabres, Victor Ricardo Candido Torres da Silvia, Siwei Zhang, Sylvia Synowiec, Silvia Honda Takada, Jubao Duan, Alexander Drobyshevsky

## Abstract

White matter injury (WMI) is a major cause of morbidity in premature infants, contributing to 5%–10% of cerebral palsy cases and up to 50% of cognitive and behavioral deficits in the United States. Two commonly used preclinical models, intermittent hypoxia (IH) and hypoxia-ischemia (HI) are widely employed to investigate the effects of WMI. The internal capsule (IC) and corpus callosum (CC) are major white matter tracts undergoing active myelination during the neonatal period, making them particularly vulnerable to hypoxic insults. This study aims to compare the effects of IH and HI models on myelination as well as the involvement of inflammatory cells in the IC and CC. We evaluated five oligodendrocyte (OL) subtypes, along with astrocytes, microglia, and activated microglia in IC and CC at postnatal day 12 (P12) and day 20 (P20) using spatial transcriptomics (CosMx, Novogene). For the HI model, C57BL/6 mice at P10 underwent permanent ligation of the left carotid artery followed by 45 minutes of hypoxia (8% O_2_ / 92% N_2_). For the IH model, P3 mice were exposed to 5% O_2_ / 95% N_2_, twice daily for five consecutive days. Animals were euthanized at P12 and P20, perfused transcardially, and brains were post-fixed in 4% paraformaldehyde, dehydrated in an ethanol series, embedded in paraffin, and coronally sectioned at 7 μm. Slides were submitted for CosMx spatial transcriptomic analysis (NanoString Technologies), and data analysis was performed using the Seurat package in RStudio. Our results demonstrate that IH and HI models affect OL populations differently, and these effects vary by brain region. In the IC, the IH model caused earlier and more pronounced changes in OL differentiation-related gene expression compared to HI. In contrast, the CC was more affected by HI. Moreover, in the HI group, mature OL s in both regions showed reduced expression of myelination-associated genes. This was accompanied by greater activation of inflammatory cells and increased intercellular communication between these cells and mature OLs, potentially contributing to the observed hypomyelination. Overall, our study provides critical insights into how each model of neonatal hypoxia differentially impacts white matter development. This knowledge can help refine preclinical strategies and guide therapeutic research tailored to the underlying pathology of each model.

## Introduction

White matter injury (WMI) is a major contributor to morbidity in premature infants, accounting for approximately 5%–10% of cerebral palsy cases and up to 50% of cognitive and behavioral impairments in the United States ^1^. In its more severe forms, WMI can progress to periventricular leukomalacia (PVL), a necrotic lesion affecting white matter near the lateral ventricles ^2^. he primary cause of WMI is typically hypoxic–ischemic injury, which compromises oxygen and blood flow to the periventricular region of the developing brain. This insult preferentially targets oligodendrocyte precursor cells (OLPCs), the key progenitors responsible for initiating myelination, leading to their apoptosis and subsequent disruption of white matter development.

Myelination is a tightly regulated, late-stage developmental process in which oligodendrocytes (OLs) generate a multilamellar myelin sheath, a specialized extension of the plasma membrane that ensheathes axons in regularly spaced segments. This sheath is essential for enabling efficient saltatory conduction of electrical impulses along nerve fibers. The process involves extensive remodeling of OL morphology and membrane architecture and is regionally and temporally heterogeneous, with the timing of myelination varying significantly across different brain regions ^3–5^.

Oligodendrocytes (OLs) are the principal myelinating cells of the central nervous system (CNS). They are essential not only for accelerating neural signal transmission through saltatory conduction but also for providing metabolic support to axons. Their precursors, oligodendrocyte progenitor cells (OLPCs), are widely distributed across the CNS and maintain the ability to proliferate and differentiate throughout life. Emerging research has revealed that, beyond their classical roles in myelin development and repair, OL lineage precursor cells (OLPCs) also contribute to motor learning and neural plasticity, indicating broader functions in CNS homeostasis and adaptability ^6^.

Numerous studies have sought to elucidate the effects of hypoxia on myelination, remyelination, and the underlying cellular and molecular mechanisms. To this end, several animal models have been developed and optimized. Among the most commonly used are the intermittent hypoxia (IH) and neonatal hypoxia–ischemia (HI) models, both of which are employed to study neurodevelopmental alterations, including deficits in myelination ^7–9^. While both models exhibit hypomyelination, they differ in pathophysiological outcomes: the IH model has been associated with hypomyelination without accompanying neuronal loss or significant neuroinflammation, whereas the HI model typically results in neuronal loss, glial activation, and a robust neuroinflammatory response.

HI model varies across studies, with induction occurring at postnatal day 1 (P1), P3, or later, depending on the specific developmental stage under investigation. Protocols often involve exposing animals to alternating hypoxic and normoxic periods over several days. The neonatal HI model is typically characterized by unilateral ischemia combined with global hypoxia, leading to variable lesion severities. These lesions are commonly classified as mild (<25%), moderate (25%–50%), or severe (>50%) based on the extent of central nervous system (CNS) damage ^10^. This model is distinguished by neuronal loss, sustained neuroinflammation that can persist for months, and hypomyelination ^11–13^. Several studies have identified multiple potential causes of hypomyelination, with increased neuroinflammation in the CNS being one of the most prominent factors ^14,15^.

Although most studies have not specifically examined whether the neonatal intermittent hypoxia (IH) model induces brain inflammation, existing evidence suggests an absence of overt cell death in this model ^7,16^. Furthermore, no significant changes in inflammatory gene expression have been reported in neonatal IH, indicating that this condition may not strongly activate classical neuroinflammatory pathways.

In contrast, the mechanisms underlying prolonged lesion development in the hypoxia-ischemia (HI) model remain incompletely understood but are believed to involve chronic inflammation and epigenetic modifications ^17^. he inflammatory response is initiated by peripheral leukocyte infiltration, activation of local microglia, and astrogliosis ^18^. Reactive astrocytes release pro-inflammatory cytokines such as tumor necrosis factor-α (TNF-α) and interleukin-6 (IL-6), which may exert sustained modulatory effects on neighboring neurons, contributing to ongoing neuroinflammation and tissue damage ^17,18^.

Hypoxia, asphyxia, and ischemia disrupt oxygen delivery and reduce metabolic support to the developing brain, leading to impaired myelination in vulnerable white matter regions during the neonatal period. Among the most affected structures are the internal capsule (IC) and corpus callosum (CC). In humans, myelination of the internal capsule begins around birth, whereas myelination in the corpus callosum can be disrupted by hypoxic events as early as 10 days postpartum^19^. Because both regions undergo active myelination during this critical developmental window, they are especially susceptible to hypoxic–ischemic injury. Additionally, oligodendrocyte progenitor cells (OLPCs) within these areas are metabolically active and highly sensitive to oxidative stress and inflammation induced by hypoxia, making them key targets of injury ^5,9^

Beyond the direct effects of neuroinflammation on oligodendrocytes and myelination, studies have identified two distinct modes of myelination: one that is independent of axonal activity and another that is activity-dependent ^20^. Leveraging the power of single-cell spatial transcriptomics (CosMx technology), this study investigated interactions between neuroinflammatory cells (astrocytes and microglia), excitatory and inhibitory neurons, and oligodendrocyte populations. We examined the transcriptomic effects of intermittent hypoxia (IH) and hypoxia-ischemia (HI) models on oligodendrocytes within key brain structures vulnerable to hypoxia, including the corpus callosum and internal capsule. Our goal was to identify key genes involved in the myelination process within each model, highlighting potential molecular targets for novel therapeutic interventions.

## 2. Materials and Method

### 2.1 Animals

For this study, we used six C57BL/6J mice (n=5/group). The mice were originally obtained from The Jackson Laboratory (Bar Harbor, Maine) and bred at the NorthShore University Health System animal facility. Pups remained with their dams, which had access to food and water ad libitum. Environmental conditions, including a 12-hour light/dark cycle and a controlled temperature of 22°C ± 2°C, were maintained. This study was approved by the Institutional Animal Care and Use Committee (IACUC) of NorthShore University Health System.

### 2.2 Experimental Groups

#### The pups were randomly assigned to the groups (Figure 1)

**Intermittent hypoxia (IH)**, and Hypoxia-ischemia (HI). For spatial transcriptomic analysis, the HI group was further divided based on hemispheric location relative to the lesion: ipsilateral (**VIP group**) and contralateral (**VCT group**) to occlusion of the left common carotid artery.

**Figure 1.**
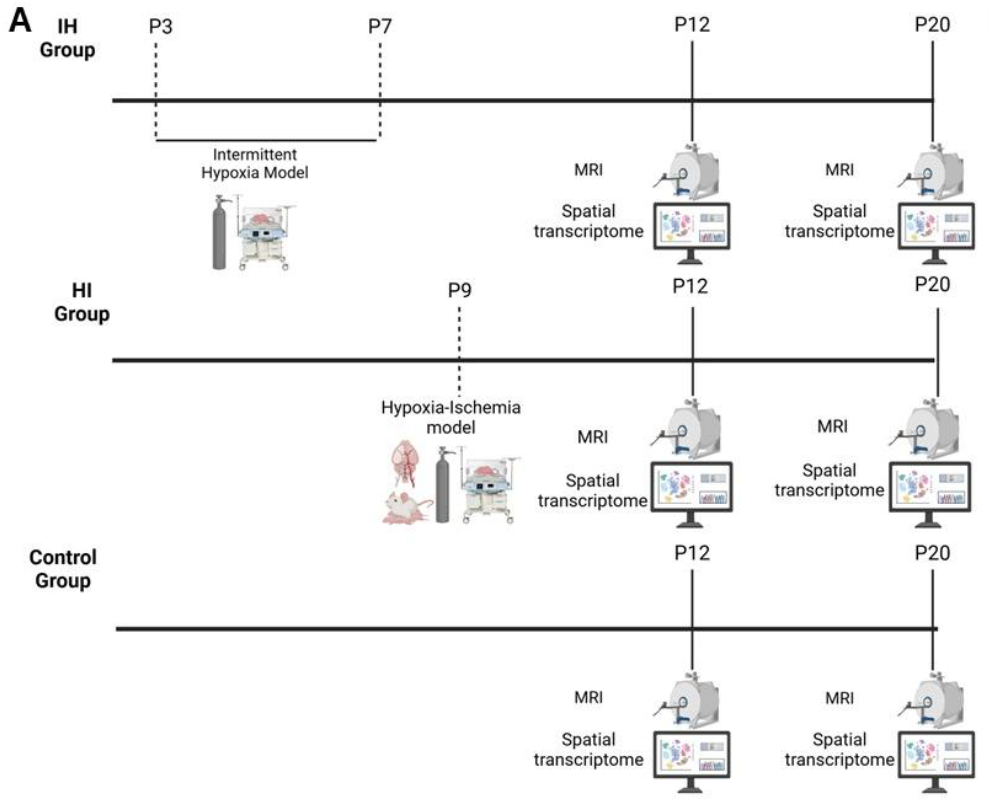
Experimental design and Image of samples used for spatial transcriptome analysis. **A**. On postnatal day 3 (P3), animals assigned to the intermittent hypoxia (IH) group were subjected to IH exposure for five consecutive days. Animals in the hypoxia-ischemia (HI) group underwent permanent occlusion of the left common carotid artery at postnatal day 9 (P9), followed by a 1–2 hour recovery period with their dams. They were then placed in a hypoxic chamber (8% O_2_ and 92% N_2_) for 45 minutes. Control animals were not subjected to either carotid occlusion or hypoxic conditions. All groups underwent MRI at postnatal day 12 (P12). One animal per group was euthanized at P12 for spatial transcriptomic analysis. At postnatal day 20 (P20), one additional animal from each group was scanned by MRI and subsequently euthanized for transcriptomic analysis.

**Figure 2.**
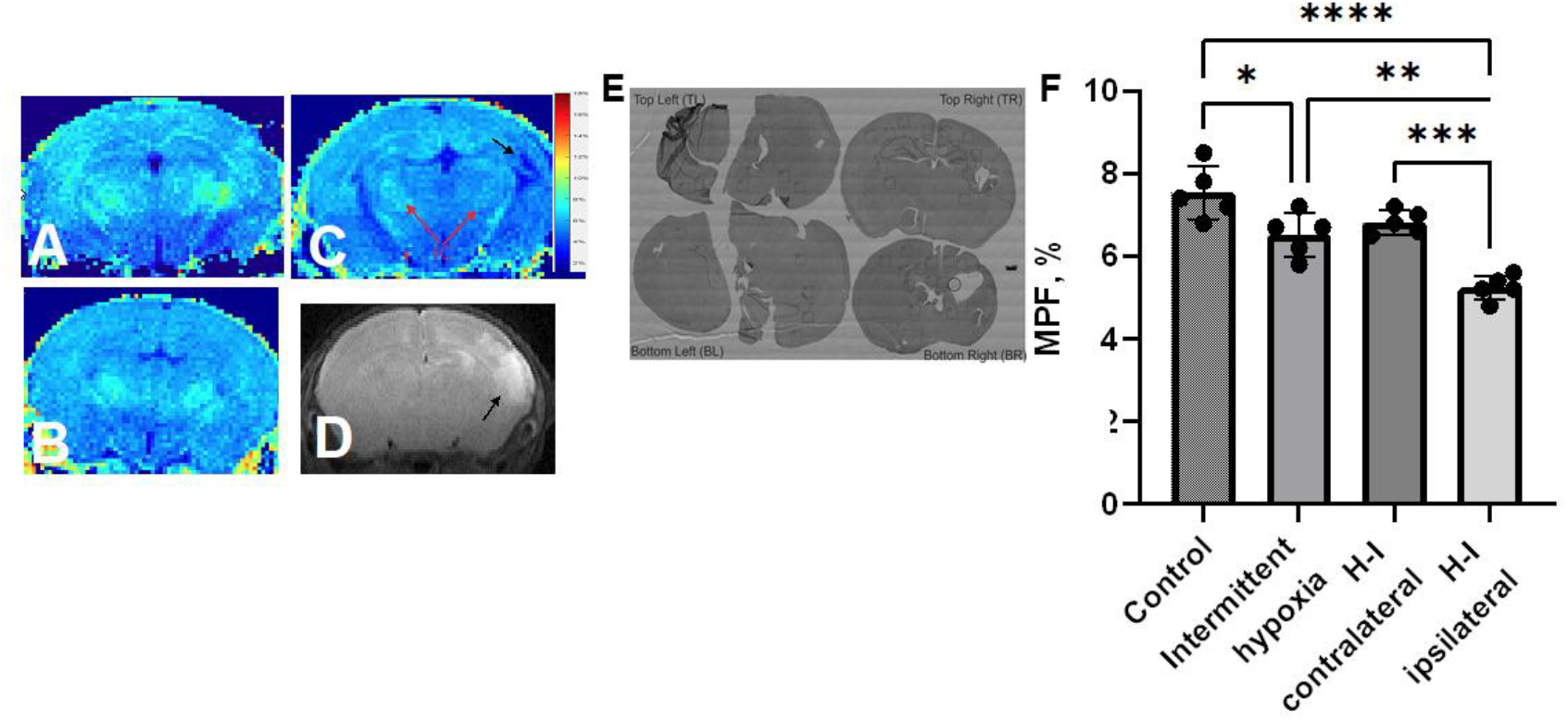
Hypomyelination in perinatal hypoxic brain injury models on myelin sensitive MRI and selection of regions of interest for special transcriptomics analysis. Macromolecular proton fraction (MPF) maps in control **(A)**, intermittent hypoxia **(B)** and hypoxia-ischemia **(C)** mouse pups at P20. Red arrows indicate internal capsule ipsi-and contralateral to H-I lesion. Pseudo color bar indicates MPF values in %. Scale bar 1 mm. Presence and extent of H-I injury was evaluated by hyperintensity on T2-weighted image (**D**, black arrow) at P12 pups, 72 hours after H-I. **E**. Paraffin sections from the corresponding MPF maps with selected regions of interest for special transcriptomics analysis. **F**. Decrease of MPF in both hypoxia models at P20, one-way ANOVA with post-hoc pairwise comparisons, *- p<0.06, **-p<0.01.

### 2.3 Intermittent Hypoxia Model

To assess the effects of intermittent hypoxia (IH), we employed a previously established protocol developed in our laboratory ^16,21^. Mouse pups were separated from their dams and placed in a 250 mL airtight chamber housed within a temperature-controlled neonatal incubator maintained at 35°C. The animals were exposed to twice-daily IH sessions, consisting of 5% oxygen (O_2_) and 95% nitrogen (N_2_). Airflow was set to 4 L/min and alternated between the hypoxic gas mixture and room air using solenoid valves operated by a programmable timer. Oxygen levels and chamber temperature were continuously monitored using calibrated sensors. The O_2_ concentration reached a nadir of 5% within 40 seconds during each hypoxic episode. The number of daily hypoxic episodes gradually decreased over the five-day exposure period as follows: 20, 18, 11, 9, and 8 episodes per day, with approximately four-hour intervals between sessions. To replicate the type of brain injury commonly observed in preterm infants, the IH protocol was initiated on postnatal day 3 (P3), a stage developmentally equivalent to that of a premature human brain, and continued through postnatal day 7 (P7), for a total of five days of intermittent hypoxia ^16^.

### 2.4 Neonatal Hypoxia-Ischemia Model

At postnatal day 9 (P9), the neonatal hypoxia-ischemia (HI) procedure was performed following the Rice-Vannucci model ^22^ and a previously established protocol from our team ^23^. Pups were anesthetized with isoflurane (5% for induction, 3% for maintenance), and a midline neck incision was made to expose the left common carotid artery, which was carefully isolated and permanently ligated using 4-0 silk suture. Following surgery, pups were allowed to recover with their dams for 1–2 hours. Subsequently, they were placed in a temperature-controlled hypoxia chamber (n = 6 per chamber) and exposed to a certified gas mixture of 8% oxygen (O_2_) and 92% nitrogen (N_2_) for 45 minutes at 37°C. After hypoxia exposure, the pups were transferred to a heated recovery box for approximately 15 minutes, then returned to their home cages with the dams.

#### In vivo MRI methods

Mice were sedated with isoflurane (Abbot, IL) inhalation, diluted in air to 5% for induction and 1.5% for maintenance. The animal’s respiration rate and rectal temperature were monitored with a small animal physiological monitor (SAII’s Small Animal Instruments, NY, USA). Body temperature was maintained at 35 C by blowing warm air from a temperature control unit. Animals were placed prone in a cradle and imaged in a 9.4 T Bruker Biospec system (Bruker Bospin, Billerica, MA). The receiver coil was a 16-mm-diameter linear surface coil (Doty Scientific, Columbia, SC). The transmitter was a 72 mm quadrature volume coil.

#### H-I injury volume on MR imaging

At P10, 24 h after H-I, MRI data were acquired in the neonatal H-I group to estimate the extent of H-I edema. T2-weighted images were acquired using a fast spin-echo RARE sequence with a TE/TR of 40/3000 ms, four signal averages, an echo train length of 8, and an in-plane resolution of 0.12 × 0.12 mm. Twenty coronal slices with a thickness of 0.7 mm were placed to cover the entire brain. To increase throughput, pups were scanned in pairs. Hyperintense areas indicating injured brain regions (edema) were manually outlined on each slice using ITK-SNAP 3.2 (itksnap.org). Volumes of edematous areas in the cortex, hippocampus, and striatum, as well as total edematous volumes, were calculated. Animals with a moderate H0I injury, characterized by an edematous volume of 5 mm^3^ and <30 mm^3^ ^24^ and at least partially preserved internal capsule, fimbria, and corpus callosum, were selected for further myelin imaging and transcriptomic analysis.

#### Macromolecular proton fraction (MPF) mapping

The degree of myelination was evaluated by MPF mapping, previously validated as an accurate in vivo measure of myelin in gray and white matter in the developing brain ^24^. 3D MPF maps were obtained from three source images (Magnetization transfer (MT)-, Proton density (PD)-, and T1-weighted) using a single-point methodology with synthetic reference images^25^ PD- and T1-weighted GRE images were acquired with TR/TE = 16/3 ms and α = 3° and 16°, respectively. MT-weighted images were acquired with a TR/TE of 22.5/3 ms and an α of 8°. The off-resonance saturation pulse was applied at the offset frequency of 4.5 kHz with an effective saturation flip angle of 500°. All images were acquired in the axial plane with whole-brain coverage and a resolution of 0.15 × 0.15 × 0.3 mm^3^. All images were obtained with four signal averages. All 3D imaging experiments employed a linear phase-encoding order with 100 dummy scans, slab-selective excitation, and a fractional (75%) k-space acquisition in the slab selection direction. To correct for field heterogeneities, 3D B0 and B1 maps were acquired using the dual-TE (TR/TE1/TE2 = 20/2.9/5.8 ms, α = 8°) and actual flip-angle imaging (AFI) (TR1/TR2/TE = 13/65/4 ms, α = 60°) methods, respectively ^26^. Imaging time was 26 min. All reconstruction procedures were performed using custom-written C-language software available at https://www.macromolecularmri.org/.

### 2.6 Sample Preparation for Spatial Transcriptomics

At postnatal days 12 (P12) and 20 (P20), animals were deeply anesthetized with isoflurane and transcardially perfused with cold 0.9% saline, followed by 4% paraformaldehyde (PFA). Brains were carefully dissected, post-fixed in 4% PFA overnight at 4°C, and then dehydrated using an automated ethanol gradient system (Leica ASP300S). Tissue samples were subsequently embedded in paraffin, coronally sectioned at 5 μm using a rotary microtome (Microm HM 340E, ThermoScientific), and mounted onto gelatin-coated slides.

To optimize spatial transcriptomic analysis, all experimental groups (IH, HI, and control) were processed on the same slide for both P12 and P20 timepoints. For the HI group, the entire brain hemisphere was included, enabling comparative analysis between the ipsilateral (VIP) and contralateral (VCT) sides relative to the lesion. In contrast, due to spatial constraints, only one hemisphere per sample was analyzed in the IH and control groups. Each brain was positioned within a standardized 20 mm × 15 mm area, ensuring consistent tissue alignment and minimizing variability in downstream processing. To facilitate anatomical orientation, an adjacent tissue section was stained with hematoxylin and eosin (H&E) to match the coordinates of the spatial transcriptomics section. Finally, two representative slides, each containing tissue from all experimental groups, were sent to NanoString for implementation of the CosMx™ Spatial Molecular Imaging (SMI) platform.

### 2.7 NanoString CosMx data analysis

#### 2.7.1 Field of view (FOV)

Differentially expressed genes (DEGs) were analyzed between experimental groups within specific brain structures using designated fields of view (FOVs). For the corpus callosum at P12, DEGs were assessed using FOV 13 (VIP), FOV 19 (VCT), and FOV 31 (IH). At P20, the corresponding FOVs were FOV 1 (VIP), FOV 7 (VCT), and FOV 25 (IH). For the fimbria, analysis at P12 included FOV 15 (VIP), FOV 21 (VCT), and FOV 33 (IH). At P20, the respective FOVs were FOV 3 (VIP), FOV 9 (VCT), and FOV 27 (IH). In the internal capsule, FOVs used at P12 were FOV 16 (VIP), FOV 22 (VCT), and FOV 34 (IH). For P20, DEGs were evaluated using FOV 4 (VIP), FOV 10 (VCT), and FOV 28 (IH).

#### 2.7.2 Cell Cluster

A critical factor in transcriptomic analysis is sample size, which in spatial transcriptomics corresponds to the **number of cells captured per field of view (FOV)**. Unlike high-throughput single-cell RNA sequencing, spatial transcriptomics is inherently limited by tissue section thickness, spatial resolution, and the size of the FOVs (as outlined in Section 2.6).

To maximize cellular representation while maintaining specificity, oligodendrocytes were classified into two major developmental clusters: immature and mature. The immature oligodendrocyte cluster included **oligodendrocyte precursor cells (OLPCs), committed oligodendrocytes (COLs)**, and **newly formed oligodendrocytes (NFOLs)**. The mature oligodendrocyte cluster encompassed **mature oligodendrocytes (MOLs)** and **myelin-forming oligodendrocytes (MFOLs)**.

#### 2.7.3 Differential gene expression analyses (DE) and Gene set enrichment analyses

Normalization was performed and differential expression of genes was statistically evaluated by using the R Bioconductor DESeq2 package (https://bioconductor.org/packages/release/bioc/html/DESeq2.html). Gene ontology (GO) term enrichment analysis for differentially expressed genes and cluster analysis were performed using the Cluster profile package and the function EnrichGO, to perform this analysis the Genome wide annotation for Mouse (org.Mm.eg.db) was used from Bioconductor (https://bioconductor.org/packages/release/data/annotation/html/org.Mm.eg.db.html). GO terms were represented according to adjusted p-value, the number of genes related to them, and their ratio in relation to the database. A pseudobulk RNA analysis was also performed for some chosen genes in the panel, using the AverageExpression function of Seurat library

#### 2.7.4 Cell-to-cell communication analyses

First, to calculate the Cell-to-cell communication, we calculated the proximity frequency between two sets of cells, i.e., Cell type A and cell type B, distances. To compute these distances, we use the information presented in the cosmix database. First the real cell proximity was calculated considering the average number of cell B counted in the radius of cell type A, the radius of proximity which we chose was of 30um(or 250 pixels, since the comix database is in pixels we have to use the conversion rate of 0.12) considering the Euclidean distance between each Cell TypeA and all Cell Type B, taking then the mean of these counts to obtain the mu_true. After that, the background model was determined by randomly redistributing the typeB cells across the image while keeping the type A cells fixed, then the mean between those cells and the standard deviation of this distribution is also computed.

Then we calculate the enrichment of type_B around type_A with the log(fold chance):

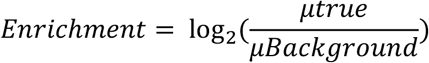

After that the Z-score was calculated to measure how different the observed proximity is from the expected background, and then a statistical test with p-value was performed. The p-value was corrected for multiple comparisons with the FDR.

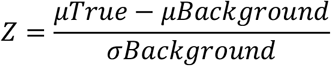

### 2.8 Statistics and Reproduction

No statistical methods were used to predetermine sample size. Data were included in the analysis unless classified as low quality, in which case cell segments or spots were filtered out as detailed in the Visium and CosMx analysis sections. Experiments were not randomized, and investigators were not blinded to group allocation during experimentation or outcome assessment. All statistical tests used are described in the main text and detailed in the Methods section below.

## Results

In this study, we evaluated RNA expression in immature and mature oligodendrocyte populations across two models of neonatal hypomyelination: Intermittent Hypoxia (IH) and Hypoxia-Ischemia (HI). The HI model was further subdivided into two groups based on the side relative to the carotid artery occlusion: Vannucci Contralateral (VCT) and Vannucci Ipsilateral (VIP). For analysis, oligodendrocyte-lineage cells were classified into two main clusters: immature and mature. The immature cluster included Oligodendrocyte Precursor Cells (OLPCs), Committed Oligodendrocytes (COL), and Newly Formed Oligodendrocytes (NFOL). The mature cluster comprised Myelin-Forming Oligodendrocytes (MFOL) and Mature Oligodendrocytes (MOL).

### Intermittent hypoxia has hard time for mature the progenitors cells in corpus callosum

The study analyzed differentially expressed genes (DEGs) among the IH, VCT, and VIP groups and identified the associated molecular mechanisms in the corpus callosum (CC) (Fig. 3). Each group exhibited gene expression profiles that may contribute to reduced differentiation of immature oligodendrocytes in the CC at P12. Notably, a comparison between the IH and VIP groups revealed alterations in pathways related to developmental cell growth (GO:0048588) and developmental growth involved in morphogenesis (GO:0060560) (Figure GO IH vs. VIP).

**Figure 3.**
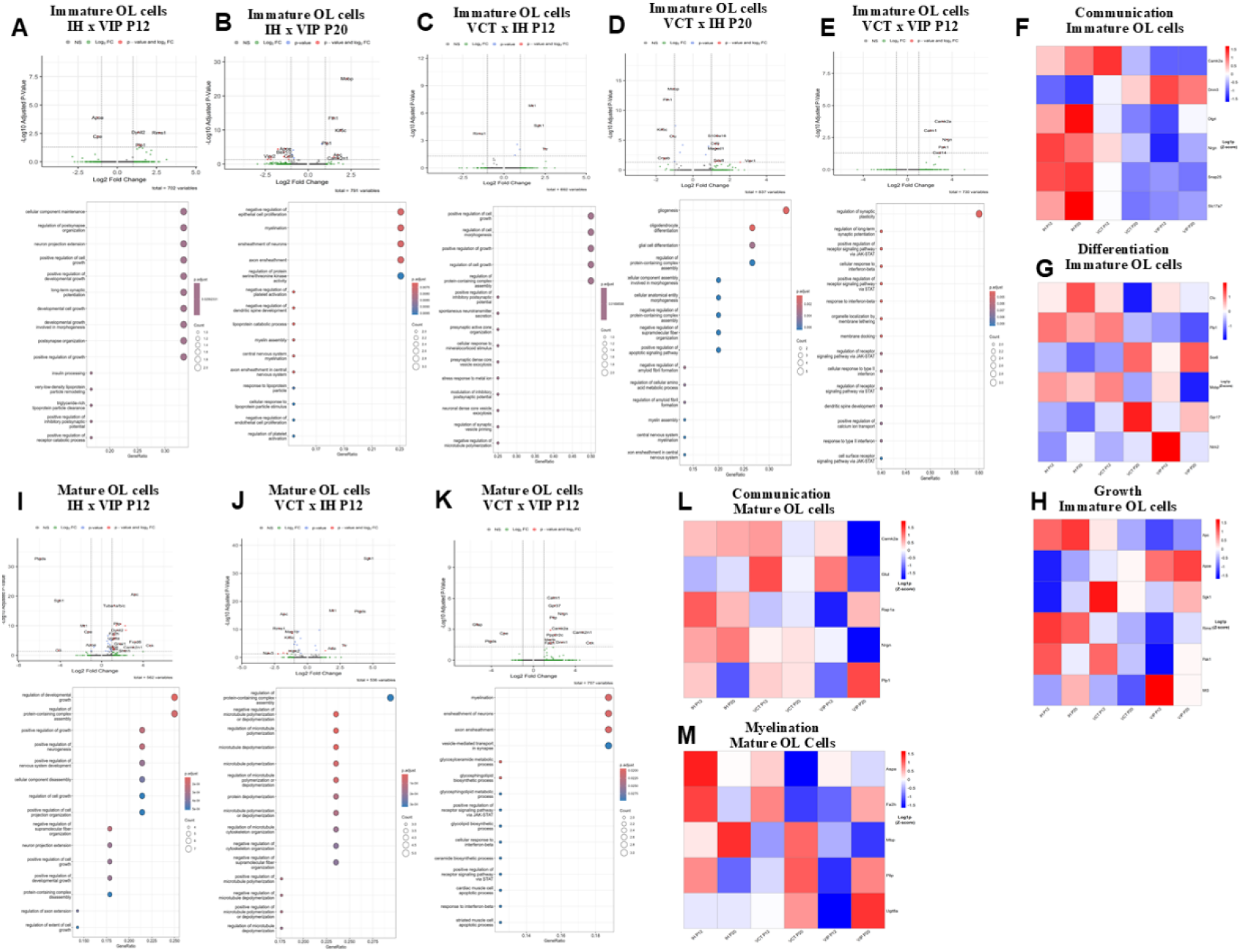
Differential gene expression and enriched biological pathways in oligodendrocyte populations. in the corpus callosum. Volcano plots show differentially expressed genes (DEGs) between the IH and VIP groups at postnatal day 12 (P12) and day 20 (P20) in immature oligodendrocytes (OLs) **(A, B)** and mature OLs **(I)**, along with enriched Gene Ontology (GO) biological processes. DEGs and GO enrichment between the VCT and IH groups at P12 and P20 are shown for immature OLs **(C, D**) and mature OLs **(J)**. Comparisons between VCT and VIP groups at P12 are presented for immature OLs **(E)** and mature OLs **(K)**. DEGs were identified using an adjusted p-value < 0.05 and log_2_ fold change (log_2_FC) > 0. Heatmaps illustrate expression patterns of genes related to structure and communication **(F)**, differentiation **(G)**, and growth **(H)** in immature OLs, as well as communication **(L)** and myelination-related genes (M) in mature OLs, across ages and experimental groups. Expression levels are represented as log_2_(RPKM).

The *Clu* gene, known to inhibit the differentiation of immature oligodendrocytes when upregulated, was increased in all three groups (IH, VIP, and VCT). Conversely, *Gpr17*, which promotes oligodendrocyte differentiation when upregulated, was downregulated across all groups. Additionally, *Plp1* and *Ntrk2* were specifically altered in the VIP group (Fig. 3. G).

At P20, gene expression analysis showed that the IH group had minimal changes, whereas the VCT and VIP groups exhibited greater alterations in genes associated with cell differentiation and growth compared to the IH group (Fig. 3. G and H). When comparing VCT and IH groups, several pathways involved in cellular growth and differentiation were significantly altered, including oligodendrocyte differentiation (GO:0048709), gliogenesis (GO:0042063), and glial cell differentiation (GO:0010001) (Fig. 3. D).

Within-group temporal analysis revealed that in the HI model overall, gene expression related to differentiation and growth remained relatively stable across timepoints. However, in the VCT group, greater changes in these genes were observed at P20 compared to P12. In contrast, the VIP group displayed an opposite pattern, with more prominent gene alterations observed at P12 than at P20.

### Alterations in communication among immature cells were more pronounced in the VIP and VCT groups compared to the IH group in the corpus callosum

Comparison of gene expression in immature oligodendrocyte cells between the IH and VIP groups at P12 revealed alterations in pathways related to the regulation of postsynapse organization (GO:0099175) and postsynapse assembly (GO:0099173) (Fig. 3A). Differentially expressed genes (DEGs) included synapse-associated genes such as *Camk2a, Dlg4, Nrgn, Snap25*, and *Slc17a7*, which were downregulated in the VIP group. Between the VCT and IH groups, pathways related to synaptic function (GO:0097151, GO:0061669, and GO:1990709) were also altered at P12. However, by P20, no significant changes in synapse-related pathways were observed between any of the groups.

When comparing VCT and VIP groups, pathways involved in the regulation of synaptic plasticity (GO:0048167), regulation of long-term synaptic potentiation (GO:1900271), and signaling via the JAK-STAT pathway were differentially regulated. Notably, the VCT group exhibited increased numbers of altered genes at P20 compared to P12, whereas the IH group maintained relatively stable gene expression across these ages. The VIP group, in contrast, showed more pronounced gene alterations at P12 than at P20 (Fig.3).

### Hypoxia-ischemia model has more difficult for express myelination genes by mature cells

In mature oligodendrocyte cells, an increase in the alteration of genes involved in myelination was observed at P20 compared to P12 in the IH group. Conversely, the VCT and VIP groups showed a decrease in the number of altered genes with advancing age (from P12 to P20). Gene-specific analysis revealed that the *MBP* gene was upregulated in the HI group at both P12 and P20, whereas in the VCT group, *MBP* expression was reduced at P12 but increased by P20. The VIP group exhibited reduced *MBP* expression at both ages.

Comparisons between the VCT and VIP groups at P12 highlighted significant alterations in pathways related to myelination (GO:0042552) and ensheathment of neurons (GO:0007272). Additionally, pathways involving JAK-STAT signaling (GO:0046427 and GO:1904894) and cellular response to interferon-beta were affected, underscoring the role of inflammation in this model. Among mature cells, calcium ion-regulated exocytosis of neurotransmitters (GO term) was altered between the VCT and IH groups, while similar pathway changes were not detected in other comparisons (Figure 3I–K). However, when analyzing genes specifically from the VIP group, a greater number of communication-related genes were found to be altered at both P12 and P20 (Figure 3L).

### Genes about differentiation of immature cells in Internal Capsule were early impacted by intermittent episodes of hypoxia

Analysis of immature cells in the internal capsule revealed that the IH group exhibited a greater number of altered genes related to differentiation and growth at P12 compared to the VCT and VIP groups (Fig. 3H and I). Additionally, pathways associated with myelination (GO:0042552) and axon ensheathment (GO:0008366) were upregulated in the VIP group relative to the IH group, driven by increased expression of genes such as *Mbp, Mobp*, and *Bcas1* (Fig. 4Q). Similar upregulation of these pathways and genes (*Mbp* and *Mobp*) was observed in the VCT group compared to IH.

**Figure 4.**
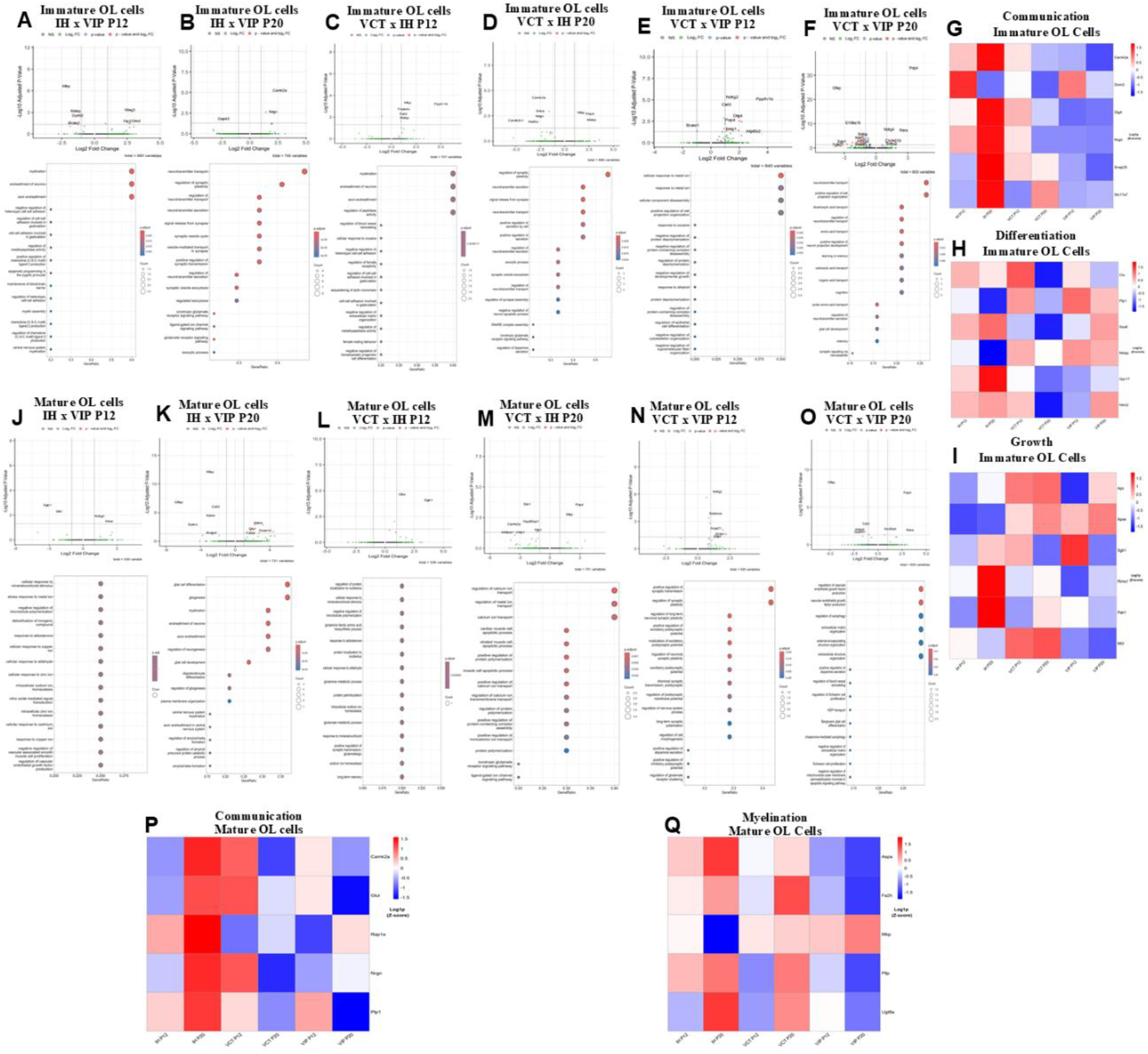
Transcriptomic analysis of immature and mature oligodendrocyte populations in the internal capsule across IH, VCT, and VIP groups. Volcano plots display differentially expressed genes (DEGs) and enriched Gene Ontology (GO) biological pathways between the IH and VIP groups at postnatal days 12 and 20 (P12 and P20) for immature oligodendrocytes (OLs) **(A, B)** and mature OLs **(J, K)**. Comparisons between the VCT and IH groups are shown at P12 and P20 for immature OLs **(C, D)** and mature OLs **(L, M)**. DEGs and GO terms between the VCT and VIP groups are illustrated for immature OLs at P12 and P20 **(E, F)** and for mature OLs **(N, O)**. Heatmaps highlight the expression of genes associated with structural organization and communication **(G)**, differentiation **(H)**, and growth **(I)** in immature OLs, as well as communication **(P)** and myelination **(Q)** genes in mature OLs, across both ages and experimental groups. Differential expression was defined as adjusted p-value < 0.05 and log_2_ fold change (log_2_FC) > 0. Gene expression levels are presented as log_2_(RPKM).

The gene *Sgk1*, which influences the oxidative stress response, was upregulated in both VCT and VIP groups at P12 but decreased by P20. Notably, the VIP group showed an increase in the number of altered genes related to differentiation and growth at P20 compared to P12, whereas the IH group exhibited fewer altered genes at P20 than at P12. The VCT group presented only three altered genes related to differentiation and growth.

Regarding communication-related genes, the VIP group displayed alterations at both P12 and P20 compared to the other groups. In contrast, the IH group had changes limited to *Snap25* and *Slc17a7* at P12, and only *Dnm3* at P20. At P20, pathways involved in ionotropic glutamate receptor signaling (GO:0035235) and glutamate receptor signaling (GO:0007215) were upregulated in the IH group compared to VIP. Furthermore, when comparing IH and VCT groups, pathways related to regulation of synaptic plasticity (GO:0048167) and synaptic signal release (GO:0099643) were upregulated in IH, supported by increased expression of genes such as *Camk2a, Snca, Snap25, Nrgn*, and *Mef2c*.

### The myelination genes were more affected by Hypoxia-Ischemia than intermittent hypoxia in Internal capsule

In mature cells, communication-related genes were downregulated in the VCT group but upregulated in the VIP and IH groups at P20. The IH group showed significant enrichment in calcium transport and glutamate signaling pathways. Myelin-related genes in VCT were downregulated at P12 but increased at P20, while in VIP, they remained mostly downregulated at both ages except for *Mbp*. In the IH group, *Ugt8a* was downregulated at P12 and *Mbp* at P20. Myelination pathways were altered at P20 in both HI and VIP groups.

In the internal capsule, IH immature cells showed upregulation of synapse-related pathways at P20, and mature cells had increased expression of postsynaptic receptor regulation genes. In VCT, immature cells had higher expression of genes involved in neurotransmitter reuptake and myelination at P12. Mature VCT cells showed increased expression of genes promoting cell growth across ages. In the VIP group, immature cells had increased expression of genes linked to cell division, gliogenesis, and sodium ion homeostasis at P20. Mature VIP cells showed upregulation of synaptic function pathways at P20.

### The cellular interactions between inflammatory cells and immature oligodendrocytes exhibit age-dependent differences between the neonatal HI and IH groups

One of the hallmark features of ischemia models is the heightened activation of neuroinflammatory cells and associated inflammatory pathways. To assess microglial activation, subpopulation analysis was conducted to distinguish resting from activated microglia (Figure 5A). Additionally, the number of cells within each brain region was quantified to better understand cellular dynamics and interactions (Figure 5B–E).

**Figure 5.**
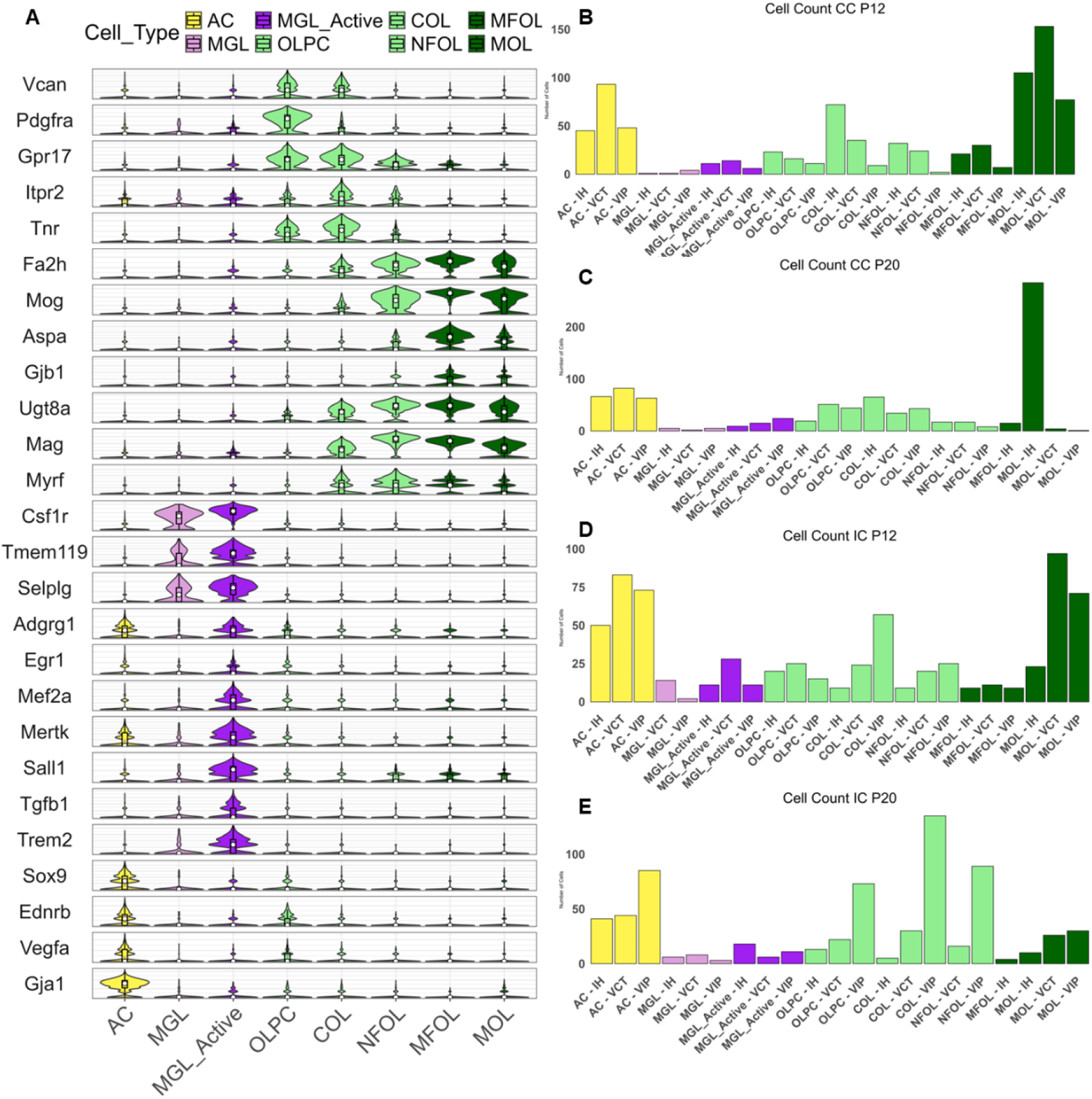
Cell identification and quantification in anatomical regions of mice at P12 and P20. Each cluster is color-coded by cell subtype, with cluster boundaries outlined and labeled by cell type. Violin plots display gene expression patterns **(A)**. Quantification of astrocytes, microglia, and oligodendrocytes in the corpus callosum at P12 **(B)** and P20 **(C)**. Cell counts in the internal capsule at P12 **(D)** and P20 **(E)**.

At P12 in the corpus callosum, the HI group exhibited increased interactions between activated microglia and both OLPCs and astrocytes, along with enhanced communication between astrocytes and immature oligodendrocyte populations (OLPCs, COLs, NFOLs) and MFOLs. In contrast, the VIP group showed stronger interactions between astrocytes and mature oligodendrocytes (MOLs) (Fig. 6. E). At P20, interaction patterns shifted: the VIP group presented increased interactions between activated microglia and OLPCs, COLs, and astrocytes, while the IH group demonstrated the highest levels of interaction between astrocytes and both immature oligodendrocytes and MOLs (Fig. 6. K).

**Figure 6.**
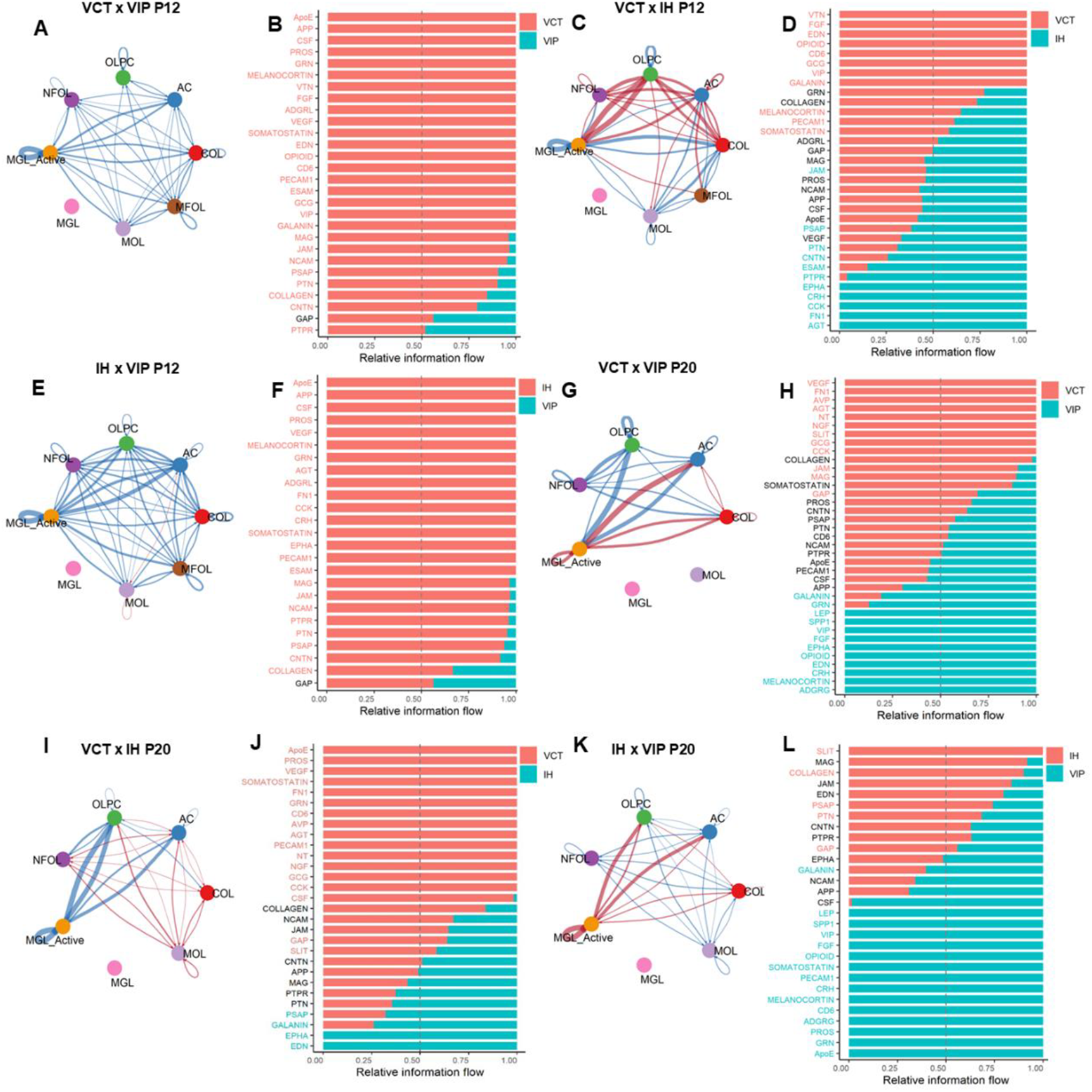
Comparison of cell-cell interactions and gene expression profiles between groups in the corpus callosum. Panels **A** and **B** show the comparison between VCT and VIP groups at P12, highlighting the most highly expressed genes in each group. Panels **C** and **D** present the comparison between VCT and IH at P12, while **E** and **F** illustrate the differences between IH and VIP at the same age. Comparisons at P20 are shown in panels G and H (VCT vs. VIP), I and J (VCT vs. IH), and **K** and **L** (IH vs. VIP), respectively

Comparing the IH and VCT groups at P12, the IH group showed more robust interactions between activated microglia and OLPCs, NFOLs, and MFOLs, and between astrocytes and OLPCs, COLs, NFOLs, and microglia. The VCT group exhibited greater interactions between activated microglia and COLs, NFOLs, and MOLs, as well as between astrocytes and MOLs (Fig. 6. C). At P20, the IH group showed increased interactions between activated microglia and OLPCs and astrocytes, and between astrocytes and all immature OL subtypes and MOLs (Fig. 6. I)

When comparing VCT and VIP groups at P12, the VCT group exhibited generally stronger cell-cell interactions across all cell types. However, neither group showed significant interactions between activated microglia and OLPCs or NFOLs. Notably, the VIP group showed increased interaction between activated microglia and COLs (Fig. 6. A).

In the internal capsule, the IH group demonstrated greater overall cellular interactions than the VIP group at both P12 and P20. When comparing VCT and IH, the VCT group showed stronger interactions at P12. At P20, OLPCs in the IH group exhibited increased interactions with activated microglia and astrocytes, while in the VCT group, astrocytes showed stronger communication with COLs and MOLs (Fig. 7. E and K).

**Figure 7.**
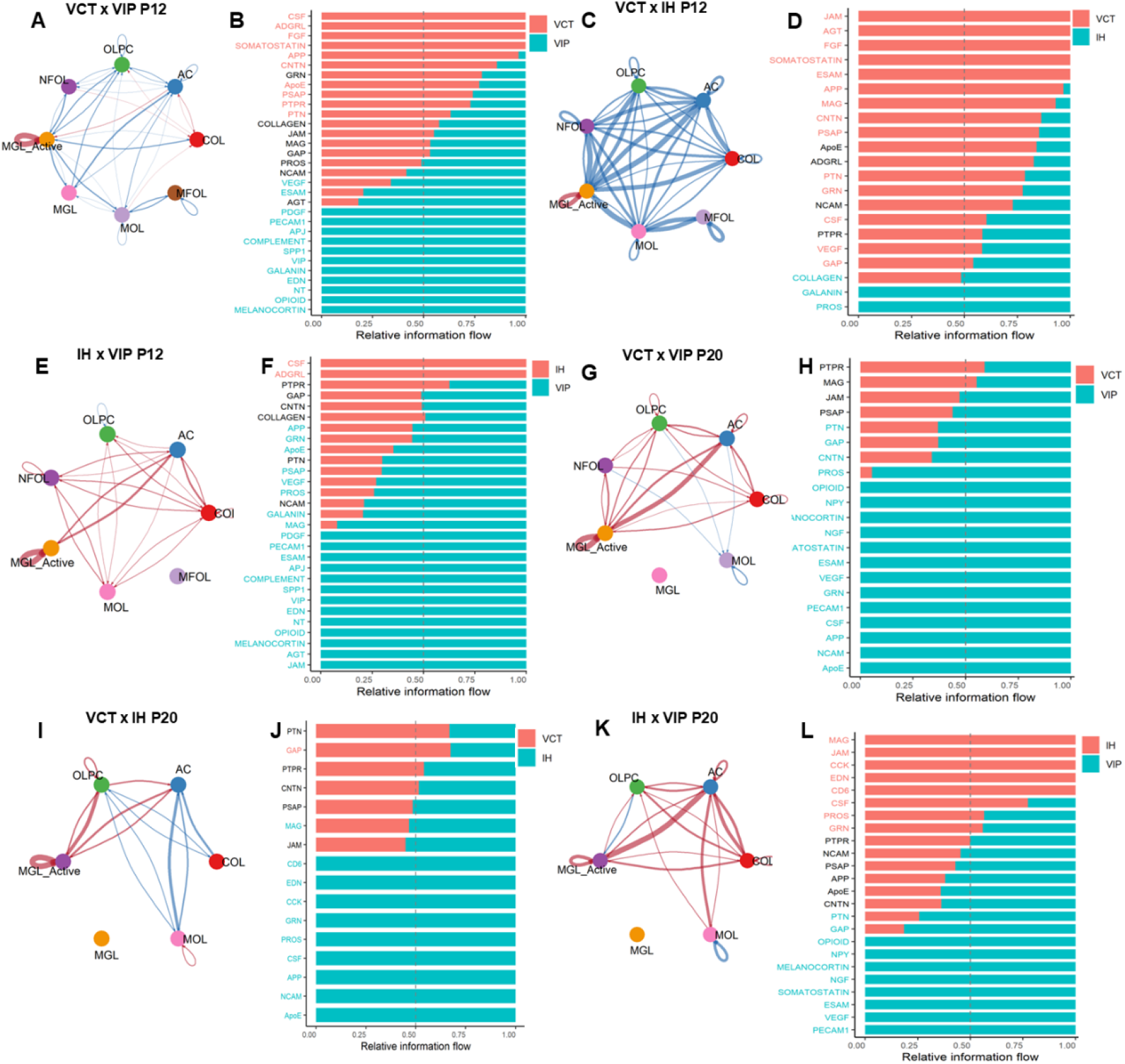
Cell–cell communication in the internal capsule. Comparison of cell–cell interactivity between groups and the most highly expressed genes in each group: VCT vs. VIP at P12 (**A** and **B**), VCT vs. IH at P12 (**C** and **D**), IH vs. VIP at P12 (**E** and **F**), VCT vs. VIP at P20 (**G** and **H**), VCT vs. IH at P20 (**I** and **J**), and IH vs. VIP at P20 (**K** and **L**).

At P12, the VCT group demonstrated greater interactions between activated microglia and multiple oligodendrocyte subtypes (OLPCs, NFOLs, COLs, MOLs, and MGLs), whereas the VIP group showed enhanced astrocyte interactions specifically with COLs (Fig. 7, A). At P20, the VIP group displayed overall stronger cellular communication than the VCT group, except for the MOL population, which in the VCT group received more signals from NFOLs, OLPCs, and astrocytes (Fig. 7. G).

## Discussion

In this study, we investigated the pathways involved in oligodendrocyte maturation, proliferation, communication, and myelination across different models of hypomyelination in two brain regions, the corpus callosum and internal capsule, at various developmental stages. MRI analysis revealed reduced myelination in both hypomyelination models compared to controls at P20 (Fig. 2F), with a more pronounced reduction observed in the group subjected to neonatal hypoxia-ischemia (HI) following left common carotid artery occlusion (VIP group).

These findings are consistent with previous studies reporting greater cellular loss and diminished myelination on the side ipsilateral to the ischemic insult^27,28^. The myelination deficit observed in the IH group is further supported by Juliano et al. (2014), who reported reduced myelination, and by Goyssakov *et al*., 2019 ^16^, who found no significant increase in cell death in mice subjected to the IH model.

In the corpus callosum, immature oligodendrocytes from animals subjected to the neonatal HI model (VCT and VIP) exhibited lower expression of genes related to communication, differentiation, and growth compared to those in the IH group at both P12 and P20. Additionally, mature oligodendrocytes showed reduced expression of genes associated with myelination and intercellular communication at P12. By P20, the HI groups also displayed a notably lower number of mature oligodendrocytes.

Myelination of the corpus callosum begins around P7^29^, becomes more prominent from P11 onward ^29,30^, and continues to increase in the number of myelinated axons through P240 ^31^. Our analysis captures two distinct stages of corpus callosum myelination: an early stage at P12, when myelination is initiating and still minimally detectable, and a later stage at P20, when myelination is more evident but still ongoing. Notably, we observed a reduced number of mature, myelinating oligodendrocytes in the neonatal HI groups.

We suggest that this reduction may be due to cell death triggered by the heightened neuroinflammatory response characteristic of the HI model ^13,17^. At P12, pathway analysis of mature oligodendrocytes revealed increased activation of inflammatory signaling, particularly the JAK-STAT pathway and the cellular response to interferon-beta, in the VIP group compared to VCT (Fig. 3K).

Interestingly, activated microglia exhibited more frequent interactions with immature oligodendrocytes in the IH group compared to the VCT and VIP groups. This suggests that in the IH model, neuroinflammation may play a greater role in modulating astrocyte–oligodendrocyte communication within the corpus callosum (Fig. 6C, E).

Analysis of pathways related to cell–cell interactions revealed increased expression of several genes associated with neuroinflammation, such as FGF, PTN, and VIP, in the VCT and VIP groups. These pro-inflammatory pathways remained elevated even at P20, supporting the hypothesis of sustained neuroinflammation in this model. Notably, these genes are known to enhance the JAK-STAT signaling pathway, and previous studies have shown that JAK3 activation is associated with increased injury severity in the neonatal HI model ^32^.

In the internal capsule, comparison between the VCT and VIP groups at P12 showed that activated microglia had stronger interactions with both immature and mature oligodendrocytes in the VCT group. In contrast, in the VIP group, the most prominent interactions involved astrocytes with COL cells and activated microglia. By P20, interaction intensity had increased globally in the VIP group, particularly involving microglia and astrocytes.

The animals were subjected to the HI model at P9. According to previous studies, phenotypical changes in microglia persisting until 17 days following injury ^33^, and astrocyte activation also can begin shortly after the insult ^13,17^. On the ipsilateral side of the common carotid artery occlusion, tissue damage is more severe. Necrotic cell death has been observed within hours of the injury ^34^, followed by a wave of apoptosis peaking within the first hours to days post-insult^27,34,35^.

These data suggest that at P12, neuroinflammatory processes are underway, particularly in the ipsilateral hemisphere. The initial loss of cells likely triggers early recruitment and activation of astrocytes, while by P20, microglial activity becomes more pronounced in this region. This sustained glial interaction may contribute to the chronic neuroinflammatory environment observed in the VIP group.

When comparing cell–cell interactions between the VIP and IH groups in the internal capsule, a greater overall level of interaction among all cell types was observed in the VIP group at both P12 and P20. In contrast, the comparison between VCT and IH groups revealed that at P12, inflammatory cells, particularly activated microglia, interacted more frequently with both mature and immature oligodendrocytes (OLs) in the IH group. By P20, the IH group continued to show strong interactions between inflammatory cells and OL subpopulations, whereas in the VCT group, interactions were more pronounced between OLPCs and NFOLs with MOLs. These findings suggest a reduction in inflammatory cell interactions in the VCT group by P20, consistent with previous reports indicating a normalization of cellular dynamics in the hemisphere contralateral to ischemia ^36,37^.

Gene expression analysis showed that in the VCT group, genes associated with oligodendrocyte differentiation were more highly expressed at P12 than at P20. In contrast, both the IH and VIP groups exhibited increased expression of these genes at P20, suggesting a delayed or prolonged differentiation process in these models. Additionally, genes related to myelination were more highly expressed in the IH group at both time points compared to the other groups.

These findings align with previous transcriptomic data from oligodendrocyte progenitor cells, which show a peak in differentiation-related gene expression at P12 followed by a decline by P80 ^5^. Other studies have reported that myelination of the internal capsule begins around P7–P10 and increases substantially between P17 and P20 ^29,30^. In our study, the VIP group displayed reduced expression of myelination-related genes at P20 compared to the other groups, suggesting impaired.

Despite the valuable findings, this study has some important limitations that should be considered. We investigated the effects of two hypomyelination models on the proliferation, differentiation, and communication of immature oligodendrocyte (OL) cells. However, due to the limited number of animals used, we were unable to assess potential sex-related differences in these processes. Additionally, the spatial transcriptomics technique used in this study has inherent limitations. Notably, it does not currently allow for the evaluation of the full transcriptome of each cell. Instead, a predefined gene panel was analyzed, which restricted the range of pathways that could be explored in this investigation.

## Conclusion

In summary, this study demonstrates that different models of hypomyelination involve distinct molecular pathways, exhibit varying gene expression profiles, and display diverse patterns of cellular interactions. Importantly, the interactions between inflammatory cells and immature oligodendrocytes vary with age between the neonatal HI and IH models. These findings enhance our understanding of potential therapeutic targets and underscore the need to identify optimal time windows for intervention. Future research should focus on developing more effective treatments, exploring sex-specific effects on oligodendrocyte maturation, and investigating additional brain regions.

## References

1. Dyet, L. E. et al. Natural History of Brain Lesions in Extremely Preterm Infants Studied With Serial Magnetic Resonance Imaging From Birth and Neurodevelopmental Assessment. Pediatrics 118, 536–548 (2006).

2. Zhu, J. et al. White matter injury detection based on preterm infant cranial ultrasound images. Front. Pediatr. 11, 1–12 (2023).

3. Marques, S. et al. Oligodendrocyte heterogeneity in the mouse juvenile and adult central nervous system. Science (80-.). 352, 1326–1329 (2016).

4. Marques, S. et al. Transcriptional Convergence of Oligodendrocyte Lineage Progenitors during Development. Dev. Cell 46, 504-517.e7 (2018).

5. Spitzer, S. O. et al. Oligodendrocyte Progenitor Cells Become Regionally Diverse and Heterogeneous with Age. Neuron 101, 459-471.e5 (2019).

6. Pukos, N., Yoseph, R. & M. McTigue, D. To Be or Not to Be: Environmental Factors that Drive Myelin Formation during Development and after CNS Trauma. Neuroglia 1, 63–90 (2018).

7. Juliano, C. et al. Mild intermittent hypoxemia in neonatal mice causes permanent neurofunctional deficit and white matter hypomyelination. Exp. Neurol. 264, 33–42 (2015).

8. Sanches, E. F., Arteni, N., Nicola, F., Aristimunha, D. & Netto, C. A. Sexual dimorphism and brain lateralization impact behavioral and histological outcomes following hypoxia-ischemia in P3 and P7 rats. Neuroscience 290, 581–593 (2015).

9. Sosunov, S. A. et al. Developmental window of vulnerability to white matter injury driven by sublethal intermittent hypoxemia. Pediatr. Res. 91, 1383–1390 (2022).

10. Fabres, R. B. et al. Therapeutic hypothermia for the treatment of neonatal hypoxia-ischemia: sex-dependent modulation of reactive astrogliosis. Metab. Brain Dis. 37, 2315–2329 (2022).

11. Netto, C. A., Sanches, E., Odorcyk, F. K., Duran-Carabali, L. E. & Weis, S. N. Sex-dependent consequences of neonatal brain hypoxia-ischemia in the rat. Journal of Neuroscience Research vol. 95 409–421 at 10.1002/jnr.23828 (2017).

12. Koo, E., Sheldon, R. A., Lee, B. S., Vexler, Z. S. & Ferriero, D. M. Effects of therapeutic hypothermia on white matter injury from murine neonatal hypoxia-ischemia. Pediatr. Res. 82, 518–526 (2017).

13. Cho, K. H. T., Davidson, J. O., Dean, J. M., Bennet, L. & Gunn, A. J. Cooling and immunomodulation for treating hypoxic-ischemic brain injury. Pediatr. Int. 62, 770–778 (2020).

14. Holloway, R. K. et al. Microglial inflammasome activation drives developmental white matter injury. Glia 69, 1268–1280 (2021).

15. Deng, Y. et al. Astrocyte-derived proinflammatory cytokines induce hypomyelination in the periventricular white matter in the hypoxic neonatal brain. PLoS One 9, (2014).

16. Goussakov, I., Synowiec, S., Yarnykh, V. & Drobyshevsky, A. Immediate and delayed decrease of long term potentiation and memory deficits after neonatal intermittent hypoxia. Int. J. Dev. Neurosci. 74, 27–37 (2019).

17. Davidson, J. O., Wassink, G., van den Heuij, L. G., Bennet, L. & Gunn, A. J. Therapeutic Hypothermia for Neonatal Hypoxic–Ischemic Encephalopathy – Where to from Here? Front. Neurol. 6, (2015).

18. Xiong, M., Yang, Y., Chen, G. & Zhou, W. Post-ischemic hypothermia for 24 h in P7 rats rescues hippocampal neuron : Association with decreased astrocyte activation and inflammatory cytokine expression. Brain Res. Bull. 79, 351–357 (2009).

19. Olivieri, B., Rampakakis, E., Gilbert, G., Fezoua, A. & Wintermark, P. Myelination may be impaired in neonates following birth asphyxia. NeuroImage Clin. 31, 102678 (2021).

20. de Faria, O., Gonsalvez, D. G., Nicholson, M. & Xiao, J. Activity-dependent central nervous system myelination throughout life. J. Neurochem. 148, 447–461 (2019).

21. Goussakov, I. et al. Abnormal local cortical functional connectivity due to interneuron dysmaturation after neonatal intermittent hypoxia. J. Neurosci. e1449242024 (2025) doi:10.1523/JNEUROSCI.1449-24.2024.

22. Rice, J. E., Vannucci, R. C. & Brierley, J. B. The influence of immaturity on hypoxic-ischemic brain damage in the rat. Ann. Neurol. 9, 131–141 (1981).

23. Drobyshevsky, A. et al. Intestinal microbiota modulates neuroinflammatory response and brain injury after neonatal hypoxia-ischemia. Gut Microbes 16, (2024).

24. Drobyshevsky, A. et al. Temporal trajectories of normal myelination and axonal development assessed by quantitative macromolecular and diffusion MRI: Ultrastructural and immunochemical validation in a rabbit model. Neuroimage 270, 119974 (2023).

25. Yarnykh, V. L. Time-efficient, high-resolution, whole brain three-dimensional macromolecular proton fraction mapping. Magn. Reson. Med. 75, 2100–2106 (2016).

26. Yarnykh, V. L. Actual flip-angle imaging in the pulsed steady state: A method for rapid three-dimensional mapping of the transmitted radiofrequency field. Magn. Reson. Med. 57, 192–200 (2007).

27. Fabres, R. B. et al. Long-Lasting Actions of Progesterone Protect the Neonatal Brain Following Hypoxia-Ischemia. Cell. Mol. Neurobiol. (/-972020) doi:10.1007/s10571-020-00827-0.

28. Skoff, R. P. et al. Hypoxic-ischemic injury results in acute disruption of myelin gene expression and death of oligodendroglial precursors in neonatal mice. Int. J. Dev. Neurosci. 19, 197–208 (2001).

29. Downes, N. & Mullins, P. The Development of Myelin in the Brain of the Juvenile Rat. Toxicol. Pathol. 42, 913–922 (2014).

30. Korrell, K. V. et al. Differential effect on myelination through abolition of activity-dependent synaptic vesicle release or reduction of overall electrical activity of selected cortical projections in the mouse. J. Anat. 235, 452–467 (2019).

31. Sturrock, R. R. Myelination of the mouse carpus callosum. Neuropathol. Appl. Neurobiol. 6, 415–420 (1980).

32. Hristova, M. et al. Inhibition of Signal Transducer and Activator of Transcription 3 (STAT3) reduces neonatal hypoxic-ischaemic brain damage. J. Neurochem. 136, 981–994 (2016).

33. Brégère, C., Schwendele, B., Radanovic, B. & Guzman, R. Microglia and Stem-Cell Mediated Neuroprotection after Neonatal Hypoxia-Ischemia. Stem Cell Rev. Reports 18, 474–522 (2022).

34. Northington, F. J., Ferriero, D. M., Graham, E. M., Traystman, R. J. & Martin, L. J. Early neurodegeneration after hypoxia-ischemia in neonatal rat is necrosis while delayed neuronal death is apoptosis. Neurobiol. Dis. 8, 207–219 (2001).

35. Shimizu, S. et al. Induction of apoptosis as well as necrosis by hypoxia and predominant prevention of apoptosis by Bcl-2 and Bcl-XL. Cancer Res. 56, 2161–2166 (1996).

36. Sejersted, Y. et al. Endonuclease VIII-like 3 (Neil3) DNA glycosylase promotes neurogenesis induced by hypoxia-ischemia. Proc. Natl. Acad. Sci. U. S. A. 108, 18802–18807 (2011).

37. Kennedy, L. et al. Lactate receptor HCAR1 regulates neurogenesis and microglia activation after neonatal hypoxia-ischemia. Elife 11, 1–21 (2022).

